# Disease-causing Slack potassium channel mutations produce opposite effects on excitability of excitatory and inhibitory neurons

**DOI:** 10.1101/2023.02.14.528229

**Authors:** Jing Wu, Imran H Quraishi, Yalan Zhang, Mark Bromwich, Leonard K Kaczmarek

## Abstract

*KCNT1* encodes the sodium-activated potassium channel Slack (KCNT1, K_Na_1.1), an important mediator of neuronal membrane excitability. Gain-of-function (GOF) mutations in humans lead cortical network hyperexcitability and seizures, as well as very severe intellectual disability. Using a mouse model of Slack GOF-associated epilepsy, we found that both excitatory and inhibitory neurons of the cerebral cortex have increased Na^+^-dependent K^+^ (K_Na_) currents and voltage-dependent sodium (Na_V_) currents. The characteristics of the increased K_Na_ currents were, however, different in the two cell types such that the intrinsic excitability of excitatory neurons was enhanced but that of inhibitory neurons was suppressed. We further showed that the expression of Na_V_ channel subunits, particularly that of Na_V_1.6, is upregulated and that the length of the axon initial segment (AIS) and of axonal Na_V_ immunostaining is increased in both neuron types. We found that the proximity of the AIS to the soma is shorter in excitatory neurons than in inhibitory neurons of the mutant animals, potentially contributing to the different effects on membrane excitability. Our study on the coordinate regulation of K_Na_ currents and the expression of Na_V_ channels may provide a new avenue for understanding and treating epilepsies and other neurological disorders.

**In brief:** In a genetic mouse model of Na^+^-activated K^+^ potassium channel gene *Slack*-related childhood epilepsy, Wu *et al*. show that a disease-causing gain-of-function (GOF) mutation *R455H* in Slack channel causes opposite effects on excitability of cortical excitatory and inhibitory neurons. In contrast to heterologous expression systems, they find that the increase in potassium current substantially alters the expression of sodium channel subunits, resulting in increased lengths of axonal initial segments.

**Highlights:** GOF mutations in Slack potassium channel cause elevated outward K^+^currents and inward voltage-dependent Na^+^ (Na_V_) currents in cortical neurons

Slack GOF does not alter the expression of Slack channel but upregulates the expression of Na_V_ channel

Slack GOF enhances the excitability of excitatory neurons but suppresses the firing of inhibitory interneurons

Slack GOF alters the length of AIS in both excitatory and inhibitory neurons

Proximity of AIS to the soma is different between excitatory neuron and inhibitory neuron

## INTRODUCTION

Epilepsy and intellectual disability (ID) are common neurological disorders with an approximate prevalence of 0.3 to 2% ^1^. Autosomal dominant pathogenic variants in the gene encoding the Na^+^ -activated K^+^ (K_Na_) channel Slack (KCNT1, K_Na_1.1) have emerged as an important cause of epilepsy and ID with a wide phenotypic spectrum, including MMPSI (malignant migrating partial seizures in infancy) and ADNFLE (autosomal dominant nocturnal frontal lobe epilepsy) ^2,3^. These variants have been shown to increase peak potassium current magnitude and produce gain-of-function (GOF), leading to network hyperexcitability and seizures ^2-5^. However, the underlying molecular mechanisms have yet to be determined.

Slack channels are highly expressed in the nervous system, including the olfactory bulb, frontal cortex, forebrain, hippocampus, thalamus, midbrain, cerebellum and brainstem ^6,7^, and play crucial roles in regulating the intrinsic excitability of neurons in multiple ways. In physiological conditions, the K_Na_ currents mediated by Slack channels can reduce neuronal excitability by activating in response to small depolarizations at the resting membrane potential ^8,9^or increase excitability by activating rapidly during an action potential (AP), which speeds up AP repolarization thereby limiting sodium channel inactivation and causing a higher AP firing frequency ^5,10-13^. The effect of a *Slack* variant on the patterns of excitability of a specific cell would therefore depend on which of these two effects take place.

Cortical networks are composed of two primary types of neurons: glutamatergic excitatory pyramidal neurons and GABAergic inhibitory interneurons, which work together to regulate many processes, including tuning the overall network excitability of the brain ^14,15^. Epilepsy has been causally linked to altered excitation/inhibition neuronal balance. Enhanced excitation in excitatory neurons and/or inhibition in inhibitory interneurons would result in hyperactivity of the epileptic circuitry ^16,17^. Two recent studies have shown that excitability and AP generation of inhibitory interneuron, but not excitatory neuron, were impaired by *Slack* GOF variants, leading to loss of inhibitory regulation and seizure susceptibility ^12,13^, suggesting that *Slack* GOF variants may have different effects on the excitability of the two specific neuron types. However, in contrast to the situation in humans, where a mutation in a single copy of the *Slack* (*Kcnt1*) gene results in serious disease, the mouse models used in these studies were homozygous for the channel mutations. Moreover, neither were the changes in K_Na_ currents characterized, nor were molecular mechanisms elucidated.

We have generated and characterized a mouse model of MMPSI that bears the *Slack-R455H* mutation (*R474H* in human numbering) ^18^. Our previous studies have demonstrated that this variant produces greatly increased K_Na_ current, together with a shift in voltage-dependence to negative potentials when expressed in heterologous cells ^3,4^. We further showed that mice bearing a homozygous *Slack-R455H* mutation are stillborn but that heterozygotes survive. These heterozygote *Slack* ^*+/R455H*^animals, however, have persistent interictal spikes, spontaneous seizures and an increased susceptibility to pentylenetetrazole-induced seizures ^18^, matching the condition for human MMPSI. In the present study, we tested whether the *Slack-R455H* GOF variant leads to network hyperexcitability and produces early-onset seizures by enhancing excitation in excitatory neurons or suppressing excitability in inhibitory interneurons. We determined that the increased K_Na_ currents have opposite effects on the excitability of the two neuron types, and that, in both cell types, the enhanced K_Na_ currents are coupled to increased expression of Na_V_ channels. Our study will provide key insights into the pathogenesis of epilepsies and associated ID in patients with Slack GOF mutations and lead to novel methods and targets for treating neurodevelopmental diseases and other neurological disorders.

## RESULTS

### Slack channels are expressed in cortical excitatory glutamatergic and inhibitory GABAergic neurons

To understand the mechanisms and consequences of *Slack* GOF mutations in neuronal excitability of specific cell types, we have generated a mouse model of Slack-associated MMPSI with the *Slack-R455H* missense mutation in exon 15 that lies in site of the RCK (regulator of potassium conductance) domain of the Slack channel using the CRISPR/Cas9 technology as described previously ^18^. For some experiments, the *Slack*^*+/R455H*^mouse line was then crossed with a GAD67-GFP mouse line that selectively expresses enhanced green fluorescent protein (EGFP) in the parvalbumin (PV)-expressing interneurons for identifying and targeting GABAergic neurons. Under epifluorescence microscopy, the green fluorescence of interneurons could be successfully observed in cultured cortical neurons on DIV 14 and in cerebral cortex of 2-month-old mice (Fig. 1A).

**Figure 1.**
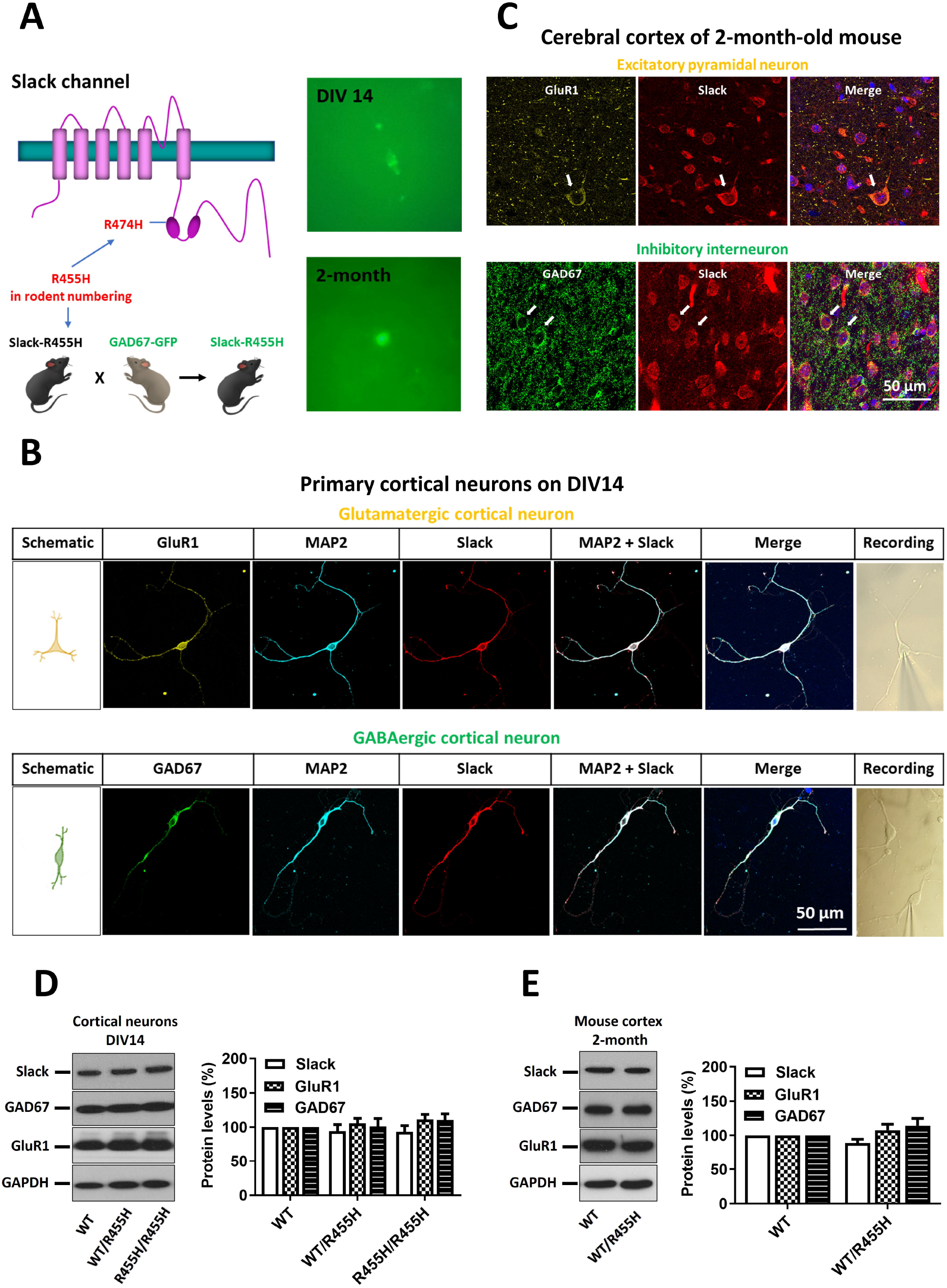
Slack channels are highly expressed in cortical glutamatergic and GABAergic neurons. **A**. *Left:* Schematic diagram of a human Slack subunit with location of *R474H residue* (*R455H* in mouse numbering). *Right:* Green fluorescence of interneurons in cultured cortical neurons on DIV 14 and in cerebral cortex of 2-month-old mouse under epifluorescence microscopy. **B**. Immunostaining of Slack channel with GluR1 (biomarker for glutamatergic neurons), GAD67 (biomarker for GABAergic neurons) and MAP2 (biomarker for soma and dendrites) in cultured cortical neurons on DIV 14. Scale bar, 50 μm. Inset shows images (40× magnification) of recorded cortical neurons under microscopy. **C**. Immunostaining of Slack channel with GluR1 and GAD67 in cerebral cortex of 2-month-old wild-type (WT) mice. Scale bar, 50 μm. **D**. Representative Western blotting and quantitative analysis of protein levels of Slack, GAD67 and GluR1 in cultured cortical neurons on DIV 14. Neuronal cultures were obtained from WT, heterozygous (WT/R455H, *Slack*^*+/R455H*^), and homozygous (R455H/R455H, *Slack*^*R455H/R455H*^) littermates. Data are shown as mean ± SEM (n = 4, one-way ANOVA). **E**. Representative Western blotting and quantitative analysis of protein levels of Slack, GAD67 and GluR1 in cerebral cortex of 2-month-old WT and *Slack*^*+/R455H*^ mice. Data are shown as mean ± SEM (n = 4, Unpaired t-test).

We first carried out immunolocalization to examine the expression of Slack channels in cultured cortical neurons. Co-staining with antibodies against GluR1 and GAD67, biomarkers for glutamatergic and GABAergic neurons, respectively, we observed that Slack channels were highly expressed in both glutamatergic and GABAergic neurons (Fig. 1B). Slack channels were expressed at the cell body and dendrites of the neurons, as indicated by its colocalization with microtubule-associated protein 2 (MAP2), a dendritic marker (Fig. 1B). Moreover, the morphological features of the glutamatergic and GABAergic neurons in primary culture were different. Consistent with previous studies ^19-21^, GluR1-positive cultured neurons had a bulbous, triangular cell body with prominent proximal dendrites and an apical dendrite arising from the apex of the cell body, while GAD67-positive interneuron had a fusiform soma with a bipolar morphology (Fig. 1B). We also carried out immunolocalization experiment on the cerebral cortex of intact mouse brain, where we also found Slack to be expressed in both glutamatergic and GABAergic neurons (Fig. 1C). In the intact brain, the expression pattern appeared slightly different from that of the cultured cells in that Slack channels were more highly expressed at the soma of the cells (Fig. 1C).

To test whether the *Slack-R455H* variant alters the expression of markers for glutamatergic and GABAergic neurons, we measured the protein levels of Slack, GluR1 and GAD67 in the cultured cortical neurons and cerebral cortex from WT and *Slack-R455H* mutant mice by Western blotting analysis. Protein lysates of cortical neurons obtained from WT, *Slack*^*+/R455H*^, or *Slack*^*R455H/R455H*^littermates (although the homozygotes do not survive birth, cultured neurons from embryos are viable) on DIV 14 showed that there were no significant differences of protein levels of Slack, GluR1 and GAD67 among these cultures (Fig. 1D). Similarly, protein levels of Slack, GluR1 and GAD67 of cerebral cortex obtained from 2-month-old WT and *Slack*^*+/R455H*^mice also showed no differences (Fig. 1E), suggesting that the *Slack-R455H* mutation did not grossly alter the expression of markers for glutamatergic and GABAergic neurons.

### The *Slack-R455H* mutation increases K_Na_ currents and voltage-dependent sodium currents (I_Na_) in both glutamatergic and GABAergic neurons

Disease-causing mutations in Slack have been shown to cause remarkable increases in K_Na_ currents in heterologous cells, with no change in levels of Slack protein in the plasma membrane ^3,4^. To determine whether K_Na_ currents were increased by the *Slack-R455H* mutation in specific types of neurons, we performed whole-cell voltage recording on cortical neurons obtained from WT, *Slack*^*+/R455H*^, or *Slack*^*R455H/R455H*^embryos at DIV 13-14. The recorded cortical neurons were classified as glutamatergic, fast-spiking (FS)-GABAergic, or non-fast-spiking (NFS)-GABAergic based on several criteria including *i)* green fluorescent label in GABAergic interneurons (Fig. 1A), *ii)* morphology ^19-21^and *iii)* post-immunolabeling for GluR1 and GAD67 (Fig. 1B) and *iv)* firing patterns ^12^. To isolate the K_Na_ currents, whole-cell voltage recordings were performed using physiological extracellular medium with 140 mM Na^+^ions and without Na^+^. The traces obtained without Na^+^were then subtracted from the traces obtained in the external Na^+^medium ^5^.

We observed that K_Na_ currents in both glutamatergic and GABAergic neurons were marked increased by the *Slack-R455H* mutation (Figs. 2A-2C) and the peak K_Na_ density was increased by as much as 2-4 fold in heterozygous neurons and 4-6 fold in homozygous neurons over that in WT neurons (Fig. 2D). Because sodium entry through voltage-dependent sodium channels is a major contributors to K_Na_ currents ^12,22-24^, we applied the sodium channel blocker tetrodotoxin (TTX) in independent experiments and comparing the reduction of K^+^currents in the different genotypes. After subtracting the traces obtained with TTX from the traces without TTX, we observed that TTX-sensitive K^+^currents in the neurons expressing *Slack-R455H* mutation were larger than in WT neurons (Figs. S1A and S1B). The difference current was increased by the *Slack-R455H* mutation, by as much as 3-fold in homozygous neurons and 2-fold in heterozygous neurons over that in WT neurons (Fig. S1C).

**Figure 2.**
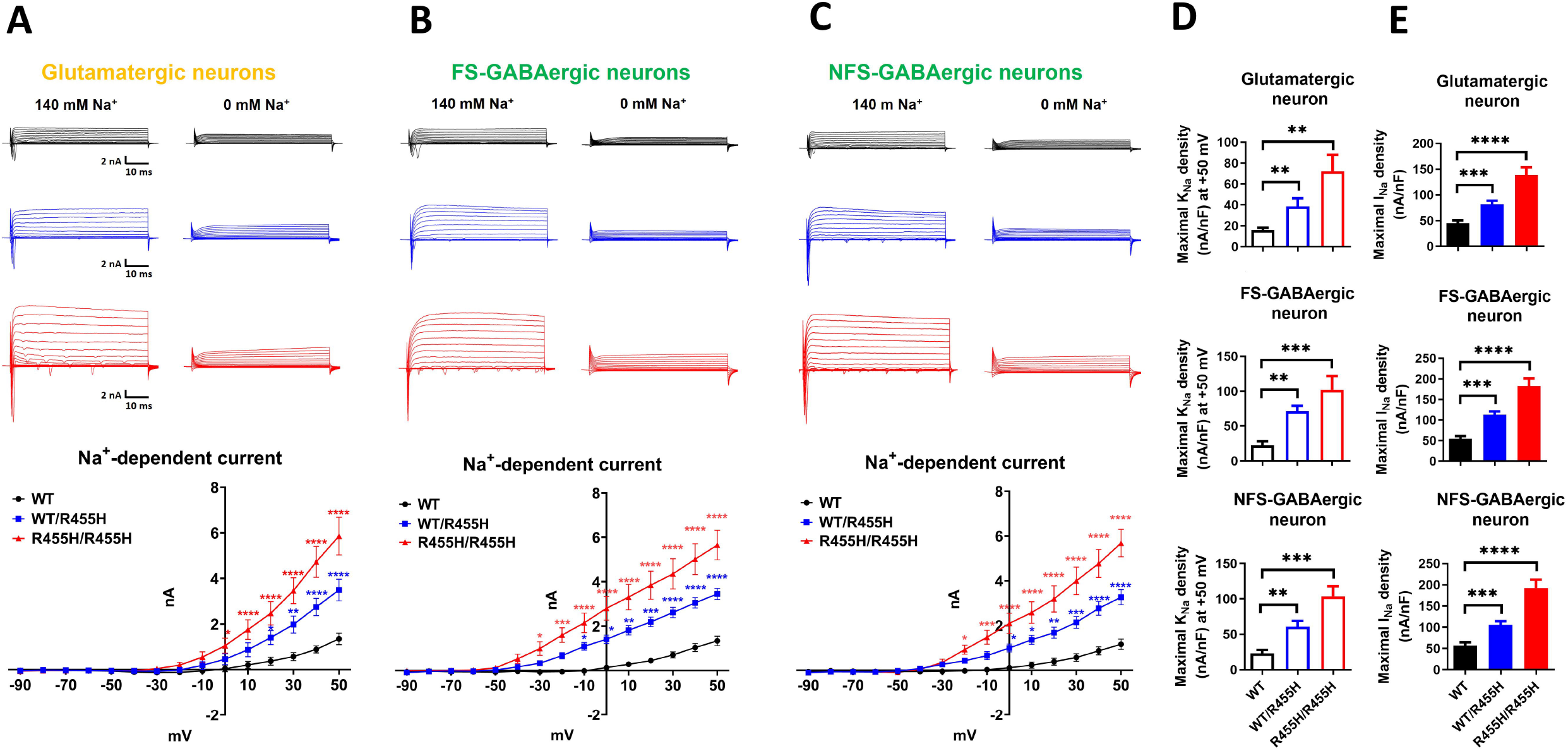
K_Na_ currents and I_Na_ are increased in both glutamatergic and GABAergic neurons expressing the *Slack-R455H* mutation. **A-C**. Representative current traces from whole cell voltage clamp recordings in physiological extracellular medium with 140 mM Na^+^ ions (*left*) and without Na^+^ ions (*right*) in response to voltage steps (−90 to +50 mV) in WT (*black*), *Slack*^*+/R455H*^ (*blue*), and *Slack*^*R455H/R455H*^ (*red*) neurons. Summary data (*bottom)* show the K_Na_ current, which was calculated by subtracting the trace obtained without Na^+^from the trace with Na^+^, for each voltage step in glutamatergic neuron (**A**), fast-spiking (FS)-GABAergic neuron (**B**), or non-fast-spiking (NFS)-GABAergic neuron (**C**). Data are shown as mean ± SEM (n = 8-12, two-way ANOVA). **D**. Maximal K_Na_ current density at +50 mV for each genotype and neuron type. Data are shown as mean ± SEM (n = 8-12, one-way ANOVA). **E**. Maximal I_Na_ density for each genotype and neuron type. Data are shown as mean ± SEM (n = 8-12, one-way ANOVA). \**p <* 0.05, \*\**p <* 0.01, \*\*\**p <* 0.001 and \*\*\*\**p <* 0.0001.

We also observed that the characteristics of the increase in K_Na_ current induced by the *Slack-R455H* mutation were different in glutamatergic and GABAergic neurons. In glutamatergic neurons, significant increases in K_Na_ current occurred in response to positive voltage steps, from +20 to +50 mV in heterozygous neurons and from 0 to +50 mV in homozygous neurons (Fig. 2A). In contrast, significant increases in K_Na_ current in FS-GABAergic neurons occurred at more negative voltage steps, from -10 mV to +50 mV in heterozygous neurons and from -30 mV to +50 mV in homozygous neurons (Fig. 2B). In the other class of interneurons, NFS-GABAergic neurons, K_Na_ currents also increased at more negative potentials, from 0 mV to +50 mV in heterozygous neurons and from -20 mV to +50 mV in homozygous neurons (Fig. 2C). Overall, the voltage-clamp recording results suggested that although the *Slack-R455H* mutation increases K_Na_ currents in all neuron types, increased K_Na_ currents at subthreshold voltages occur selectively in GABAergic neurons.

A surprising result was that, in addition to increased peak K_Na_ current density (Fig. 2D), we also found that peak I_Na_ density was increased in all neuron types expressing the *Slack-R455H* mutation, by 2-fold in heterozygous neurons and 3-fold in homozygous neurons over that in WT neurons (Fig. 2E). The results indicated that not only K_Na_ currents but also sodium currents in cortical neurons are altered by the *Slack-R455H* mutation.

### The *Slack-R455H* mutation enhances the excitability of excitatory neurons but suppresses inhibitory interneurons

To investigate further the effects of *Slack-R455H* mutation GOF on neuronal excitability of specific neuron types, we carried out whole-cell current recording on the cortical neurons from WT, *Slack*^*+/R455H*^, or *Slack*^*R455H/R455H*^embryos at DIV13-14. The electrophysiological properties of neurons were determined by injecting 200 ms square current pulses incrementing in 20 pA steps, starting at -20 pA. To compare the maximum number of action potentials (APs), 1.5 s square current pulses in 20 pA steps were injected until the number of APs per stimulus reached a plateau phase.

First, we compared the firing properties of glutamatergic neurons from different genotypes. We observed that AP half width was decreased while afterhyperpolarization (AHP) were increased in the neurons expressing *Slack-R455H* mutation when compared with WT neurons (Figs. 3A, 3C and 3D). However, rheobase, input resistance, and resting membrane potential showed no significant differences among all three genotypes (Figs. 3E, 3F and S2A). Other electrophysiological parameters, including AP amplitude, AP depolarization rate and repolarization rate, were raised by the *Slack-R455H* variant in glutamatergic neurons (Figs. 3B, S2C and S2D). Furthermore, the frequency of APs evoked by 1.5 s depolarizing currents was greatly increased in the *Slack-R455H* mutant neurons relative to that of WT neurons (Figs. 3G, H). Significant increases in AP frequency occurred from 240 pA to 260 pA in heterozygous neurons and from 160 pA to 280 pA in homozygous neurons (Fig. 3H). In addition, the maximum number of APs that could be evoked in *Slack*^*455H/R455H*^neurons was twice that in WT neurons while *Slack*^*+/R455H*^neurons were capable of firing at a rate 1.5-fold over that in WT neurons (Fig. 3I).

**Figure 3.**
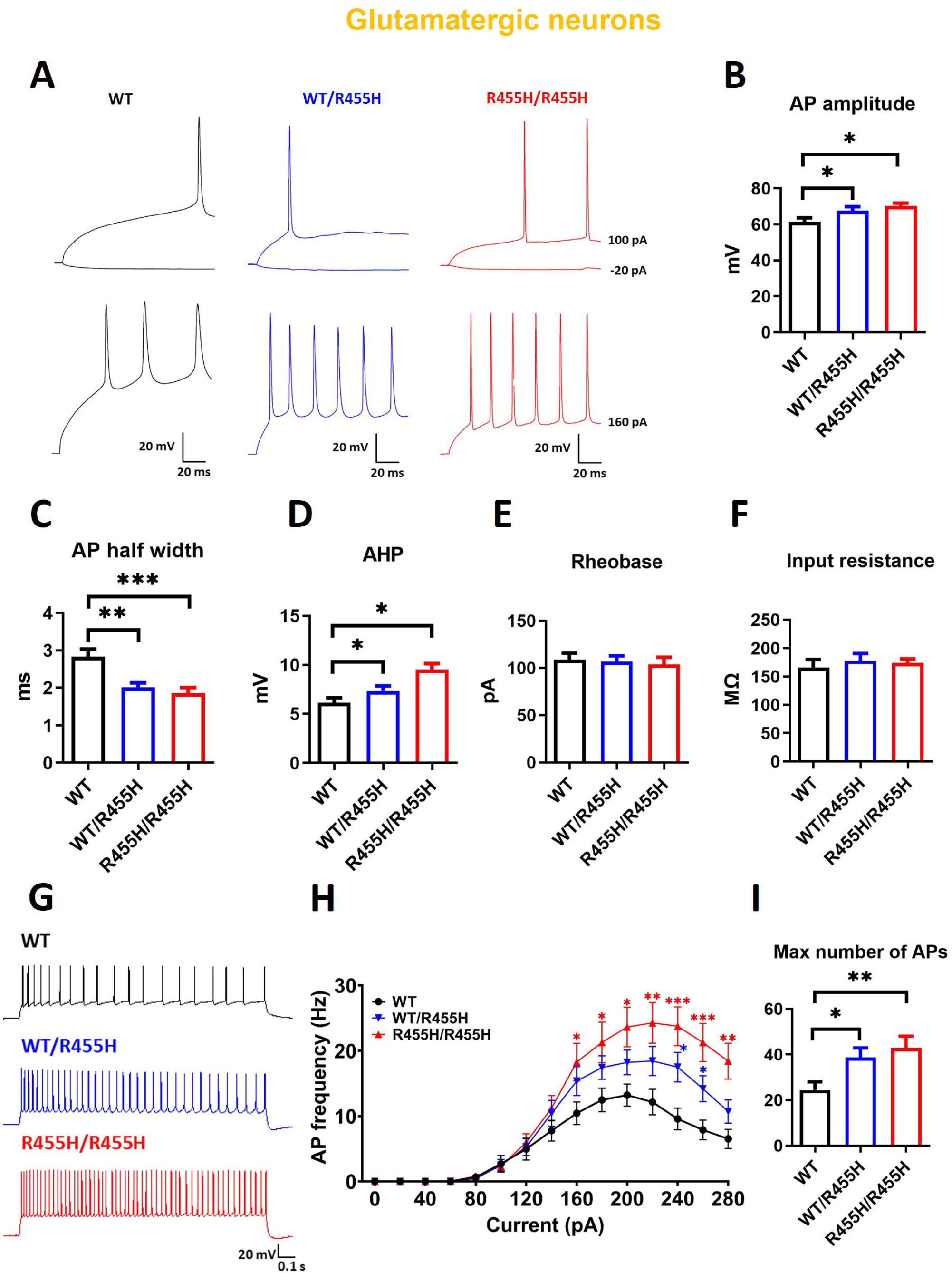
The *Slack-R455H* mutation enhances the excitability of glutamatergic neurons. **A**. Representative traces from whole cell current clamp recordings from glutamatergic neurons in response to current step injections from WT (*black*), *Slack*^*+/R455H*^ (*blue*), and *Slack*^*R455H/R455H*^ (*red*) littermates. To compare the electrophysiological properties, neurons were injected with 200 ms square current pulses incrementing in 20 pA steps, starting at -20 pA. To compare the maximum number of APs, 1.5 s square current pulses in 20 pA steps were injected until the number of APs per stimulus reached a plateau phase. AP amplitude (**B**), AP half width (**C**), AHP (**D**), Rheobase (**E**), and input resistance (**F**) of recorded neurons from each genotype. Data are shown as mean ± SEM (n = 15-20, one-way ANOVA). **G-H**. Example traces and summary data showing the number of APs per current injection step in WT (*black*), *Slack*^*+/R455H*^ (*blue*), and *Slack*^*R455H/R455H*^ (*red*) neurons. Data are shown as mean ± SEM (n = 12-20, two-way ANOVA). **I**. Maximal number of APs for each genotype. Data are shown as mean ± SEM (n = 15-20, one-way ANOVA). \**p <* 0.05, \*\**p <* 0.01, \*\*\**p <* 0.001 and \*\*\*\**p <* 0.0001.

We next compared the firing properties of GABAergic neurons from the different genotypes. In contrast to glutamatergic neurons, there were no changes in AP half width or AHP in FS-GABAergic neurons among all three genotypes (Figs. 4A, 4C and 4D). The input resistance was, however, decreased while the rheobase was increased in both the *Slack*^*+/R455H*^heterozygous and *Slack*^*R455H/R455H*^homozygous neurons when compared with WT neurons (Figs. 4E and 4F). Moreover, these neurons had a more hyperpolarized resting membrane potential and AP threshold (Figs. S2E and S2F). Again, in contrast to the excitatory neurons, the frequency of APs evoked by 1.5 s depolarizing currents was decreased in the *Slack-R455H* mutant neurons relative to that of WT neurons (Figs. 4G-4I). Significant decreases in AP frequency occurred from 180 pA to 200 pA and from 360 pA to 400 pA in heterozygous neurons and from 180 pA to 260 pA and from 340 pA to 400 pA in homozygous neurons (Fig. 4H). The other class of inhibitory neurons, the NFS-GABAergic neurons expressing *Slack-R455H* mutation also showed similar alterations in rheobase, input resistance, resting membrane potential, AP threshold and firing rate to those of the FS-GABAergic neurons (Figs. 5E-5I, S2I and S2J). In addition, some other parameters, including AP amplitude, AP half width and depolarization rate, showed changes in the opposite direction to those in the glutamatergic neurons (Figs. 5B-5C, S2K).

**Figure 4.**
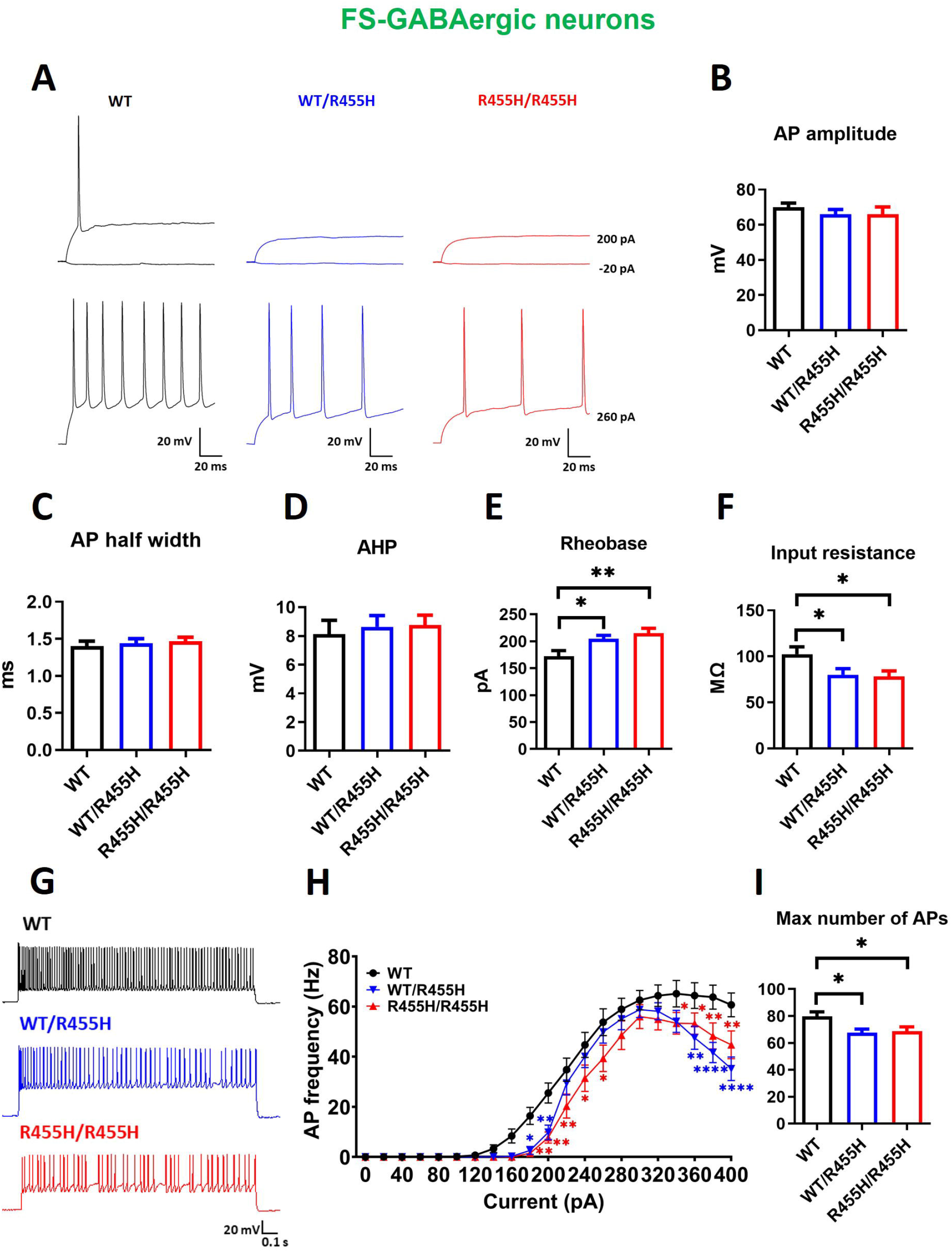
The *Slack-R455H* mutation suppresses the excitability of FS-GABAergic neuron. **A**. Representative traces from whole cell current clamp recordings from FS-GABAergic neurons in response to current step injections from WT (*black*), *Slack*^*+/R455H*^ (*blue*), and *Slack*^*R455H/R455H*^ (*red*) littermates. AP amplitude (**B**), AP half width (**C**), AHP (**D**), Rheobase (**E**), and input resistance (**F**) of recorded neurons from each genotype. Data are shown as mean ± SEM (n = 12-16, one-way ANOVA). **G-H**. Example traces and summary data showing the number of APs per current injection step in WT (black), *Slack*^*+/R455H*^ (blue), and *Slack*^*R455H/R455H*^ (red) neurons. Data are shown as mean ± SEM (n = 12-16, two-way ANOVA). **I**. Maximal number of APs for each genotype. Data are shown as mean ± SEM (n = 12-16, one-way ANOVA). \**p <* 0.05, \*\**p <* 0.01, \*\*\**p <* 0.001 and \*\*\*\**p <* 0.0001.

**Figure 5.**
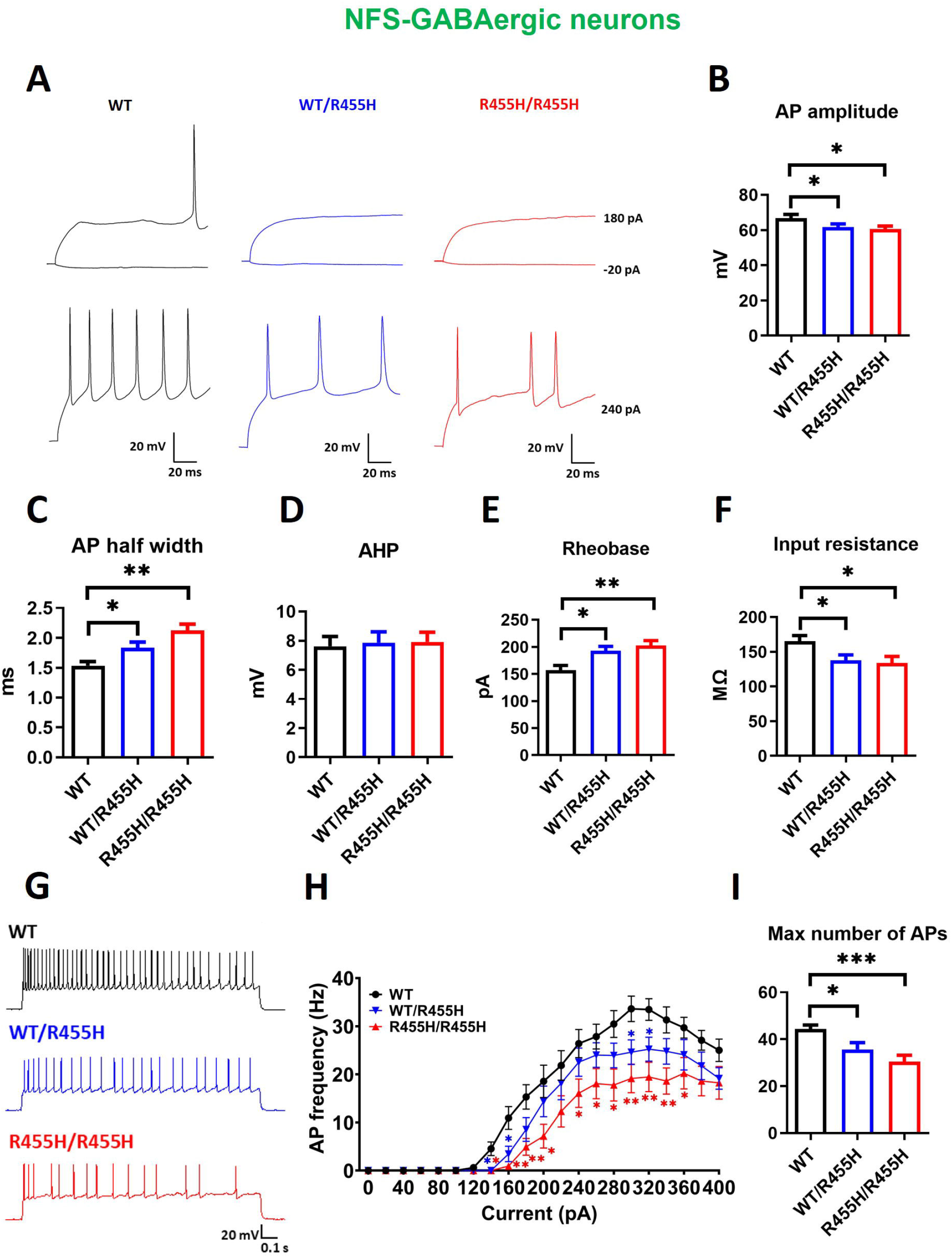
The *Slack-R455H* mutation suppresses the excitability of NFS-GABAergic neuron. **A**. Representative traces from whole cell current clamp recordings from NFS-GABAergic neurons in response to current step injections from WT (black), *Slack*^*+/R455H*^ (blue), and *Slack*^*R455H/R455H*^ (red) littermates. AP amplitude (**B**), AP half width (**C**), AHP (**D**), Rheobase (**E**), and input resistance (**F**) of recorded neurons from each genotype. Data are shown as mean ± SEM (n = 13-16, one-way ANOVA). **G-H**. Example traces and summary data showing the number of APs per current injection step in WT (black), *Slack*^*+/R455H*^ (blue), and *Slack*^*R455H/R455H*^ (red) neurons. Data are shown as mean ± SEM (n = 13-16, two-way ANOVA). **I**. Maximal number of APs for each genotype. Data are shown as mean ± SEM (n = 13-16, one-way ANOVA). \**p <* 0.05, \*\**p <* 0.01, \*\*\**p <* 0.001 and \*\*\*\**p <* 0.0001.

Overall, the current-clamp recording results indicate that the *Slack-R455H* mutation enhances the excitability of excitatory neurons but suppresses that of inhibitory interneurons.

### The *Slack-R455H* mutation upregulates the expression of Na_V_ channel subunits

To investigate the mechanism underlying the increased sodium currents in neurons from mice bearing the *Slack-R455H* mutation, we measured the protein levels of Na_V_1.1, Na_V_1.2 and Na_V_1.6, three major sodium channel isoforms in the central nervous system, in cultured cortical neurons and cerebral cortex. Western blotting analysis of cell lysates from DIV 14 cortical neurons showed that protein levels of Na_V_1.1, Na_V_1.2 and Na_V_1.6 were all upregulated in the *Slack*^*+/R455H*^heterozygous neurons and *Slack*^*R455H/R455H*^homozygous neurons when compared with WT neurons (Fig. 6A). In the cerebral cortex of 2-month-old *Slack*^*+/R455H*^mice, only the Na_V_1.6 subunit level was significantly upregulated (Fig. 6B). These results suggest that the upregulation of Na_V_ subunits in the *Slack-R455H* mutant neurons contribute to increased Na^+^influx and elevated I_Na_ and K_Na_ currents.

**Figure 6.**
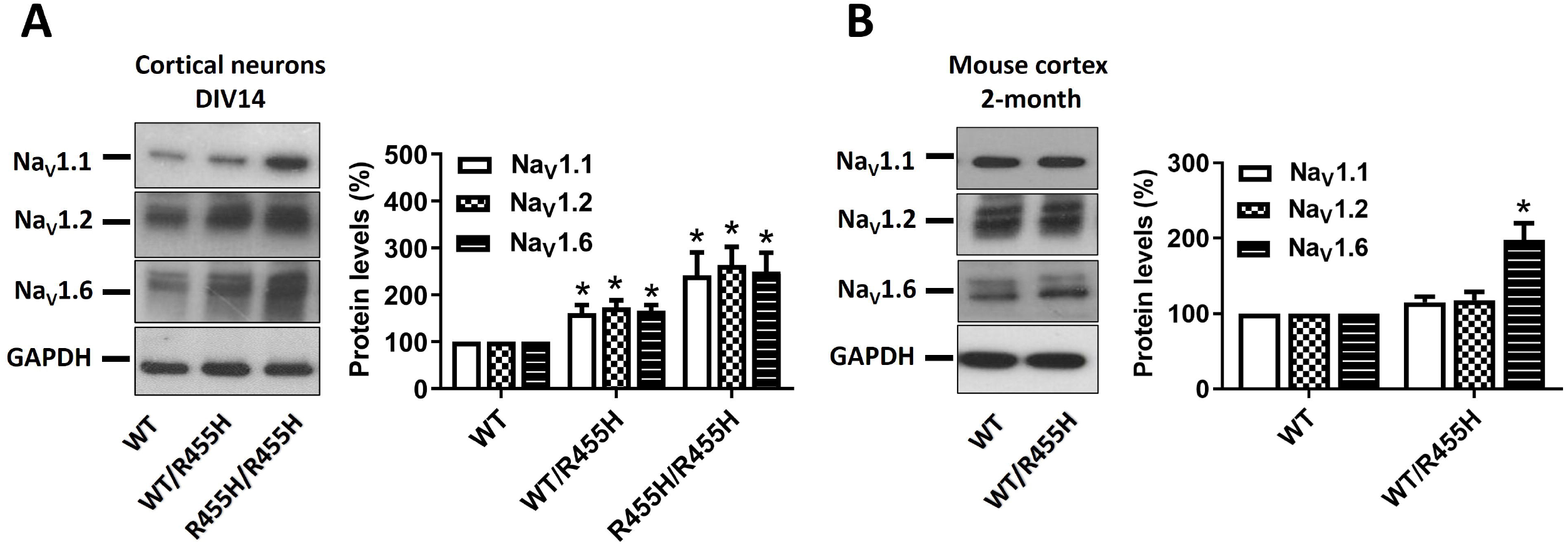
The *Slack-R455H* mutation upregulates Na_V_ channel subunit(s) expression. **A**. Representative Western blotting and quantitative analysis of protein levels of Na_V_1.1, Na_V_1.2 and Na_V_1.6 in cultured cortical neurons on DIV 14 obtained from WT, *Slack*^*+/R455H*^and *Slack*^*R455H/R455H*^littermates. Data are shown as mean ± SEM (n = 4, one-way ANOVA). **B**. Representative Western blotting and quantitative analysis of protein levels of Na_V_1.1, Na_V_1.2 and Na_V_1.6 in cerebral cortex of 2-month-old WT and *Slack*^*+/R455H*^mice. Data are shown as mean ± SEM (n = 4, Unpaired t-test). \**p <* 0.05.

### Slack channels and Na_V_ channel subunits distribute in distinct neuronal compartments in cerebral cortex

To investigate the potential cellular and molecular mechanisms by which increased K_Na_ currents have opposite effects on excitability of excitatory and inhibitory cortical neurons, we carried out co-immunolocalization experiments to determine the subcellular localizations of Slack channels and Na_V_ channels both *in vitro* and *in vivo*. We first used isoform-specific antibodies to determine the distribution of native Na_V_1.1, Na_V_1.2 and Na_V_1.6 in cultured neurons from wild type animals by immunofluorescent staining. We identified the axon initial segment (AIS) and dendrites using antibodies against ankyrin G (AnkG) and MAP2, respectively. We found that in DIV 14 cultures, Na_V_1.1 immunoreactivity co-localized with Slack in the AIS as well as the soma and dendrites of both glutamatergic and GABAergic neurons (Fig. S3A). In contrast, and consistent with previous reports ^25-28^, Na_V_1.2 (Fig. S3B) and Na_V_1.6 subunits (Fig. 7A) were highly enriched at the AIS of glutamatergic and GABAergic neurons, where they may co-localize with Slack channels.

**Figure 7.**
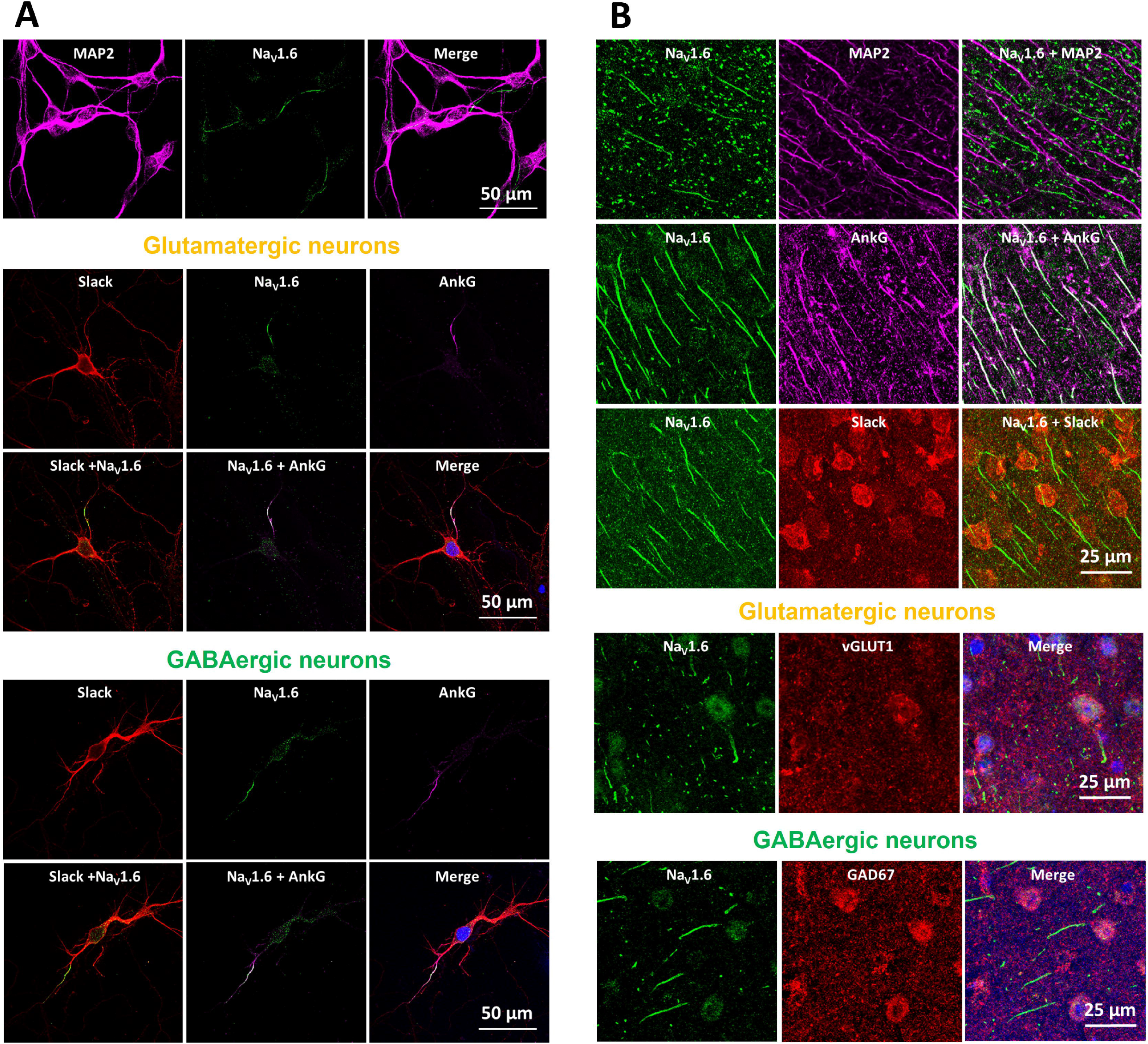
Subcellular localizations of Slack and Na_V_1.6 channels. **A**. Immunostaining of Slack and Na_V_1.6 with dendrite (MAP2) and AIS (AnkG) markers in cultured cortical neurons on DIV 14. Glutamatergic and GABAergic neurons were differentiated according to their morphological features. Scale bar, 50 μm. **B**. Immunostaining of Slack and Na_V_1.6 with MAP2 and AnkG in cerebral cortex of 2-month-old mice (top). Glutamatergic (middle) and GABAergic (bottom) neurons were differentiated by co-immunostaining with vGLUT1 and GAD67. Scale bar, 25 μm.

To characterize the cell type-specific expression and subcellular localizations of Na_V_1.1, Na_V_1.2 and Na_V_1.6 isoforms *in vivo*, immunofluorescent staining was also carried out on the frontal cortex of 2-month mice. By co-staining with markers for GABAergic neurons (GAD67) and glutamatergic neuron (vGLUT1), we were able to confirm the localization of Na_V_1.6 at the AIS of both glutamatergic and GABAergic neurons in mouse cerebral cortex (Fig. 7B), similar to our observations in cultured cortical neurons (Fig. 7A). However, Na_V_1.1 staining was specifically enriched at the AIS of GABAergic interneurons (Fig. S4A) while Na_V_1.2 staining was enriched at the AIS of glutamatergic pyramidal neurons (Fig. S4B), consistent with previous reports ^27,28^. Using a previously validated anti-Slack cytoplasmic, C-terminal antibody, we found that Slack staining was highly expressed at the soma of cerebral cortical cells (Fig. 8B), although in some cases the channels were also present at the somatodendritic region and the proximal AIS (Figs. 7B and S4). These results suggested that distribution of Slack channels and Na_V_ subunits is more restricted in adult cerebral cortex.

**Figure 8.**
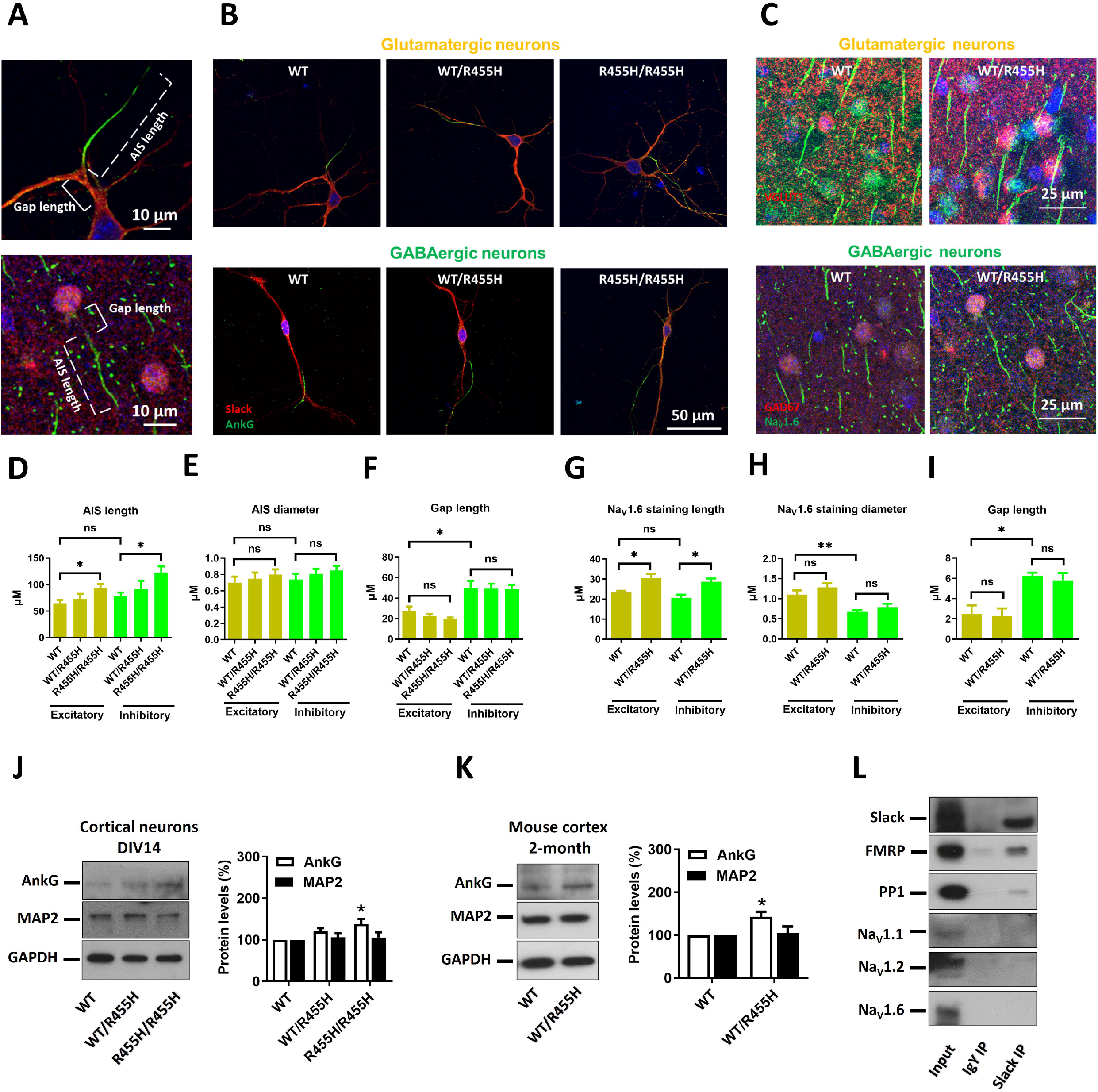
The *Slack-R455H* mutation alters the AIS length of both glutamatergic and GABAergic neurons. **A**. Representative high magnification images of a single neuron for quantification of AIS length, AIS diameter and gap length. Dotted line: AIS length; solid line: Gap length. Scale bar, 10 μm. **B**. Representative images of a single neuron and its AIS (green, AnkG) of cortical glutamatergic neurons (*top*) and GABAergic neurons (*bottom*) on DIV14. Scale bar, 50 μm. **C**. Representative images of a single neuron and its AIS (*green*, Nav1.6 staining) of pyramidal cells (*red*, vGLUT1) and interneurons (*red*, GAD67) from the cerebral cortex of 2-month-old mice. Scale bar, 25 μm. **D-F**. Quantification of overall AIS length (**D**), AIS diameter (**E**) and gap length (**F**) in excitatory glutamatergic neurons (*yellow*) and inhibitory GABAergic neurons (*green*). Data are shown as mean ± SEM (n = 3 cultures, one-way ANOVA). **G-I**. Quantification of overall AIS length (**G**), AIS diameter (**H**) and gap length (**I**) in excitatory pyramidal (*yellow*) and interneuron (*green*) populations. Data are shown as mean ± SEM (n = 3 mice, Unpaired t-test). **J**. Representative Western blotting and quantitative analysis of protein levels of AnkG and MAP2 in cultured cortical neurons on DIV14. Data are shown as mean ± SEM (n = 4, one-way ANOVA). **K**. Representative Western blotting and quantitative analysis of protein levels of AnkG and MAP2 in cerebral cortex of 2-month-old mice. Data are shown as mean ± SEM (n = 4, Unpaired t-test). **L**. Co-immunoprecipitation of Slack and Na_V_ channels from mouse cerebral cortex. Brain lysates were subjected to immunoprecipitation using anti-Slack antibody or chicken IgY, followed by Western blotting with Slack, FMRP, PP1, Na_V_1.1, Na_V_1.2, and Na_V_1.6 antibodies. \**p <* 0.05.

Collectively, the co-immunolocalization experiments reveal differential compartment-specific localization patterns of K_Na_ channels and Na_V_ channels in cultured cortical neurons and mouse cortex. The *in vitro* data shows that Slack channels are primarily localized at the soma and dendrites of cortical neurons, but also expressed at the axon where they can be co-localized with Na_V_ channels (especially with the Na_V_1.2 and Na_V_1.6 subunits). The *in vivo* data showed that Slack channels were highly expressed at the soma of cerebral cortical cells while Na_V_ channels were mainly localized at AIS. In some cases, these channels may colocalize at the proximal region of the AIS.

### The *Slack-R455H* mutation alters the AIS length of both glutamatergic and GABAergic neurons

The increase in expression of Na_V_ subunits and Na^+^current density induced by the *Slack-R455H* mutation might be expected to be reflected in the morphology of AIS. To test this possibility, we quantified the length of the AIS, determined as the length of AnkG immunostaining as well as the diameter of the AIS in wild type and mutant mice. In DIV 14 cultures (Figs. 8A and 8B), in which Na_V_1.2 and Na_V_1.6 channels are abundant at the AIS, we observed that AIS length was significantly increased in both excitatory and inhibitory neurons expressing *Slack-R455H* mutant channels when compared to that of neurons with wild type channels (Fig. 8D). When comparing all GABAergic neurons with all glutamatergic neurons, we found that there were no significant differences in AIS length and AIS diameter between these two neuron types (Figs. 8D and 8E).

We also made similar measurements in the cerebral cortices of 2-month-old mice (Figs. 8A and 8C). As in the cultured cells, we found that the length of the AIS was significantly increased in both pyramidal cells and inhibitory interneurons of mice expressing the heterozygous *Slack-R455H* mutation (Fig. 8G). Although no significant difference in AIS length was observed when comparing pyramidal cells and inhibitory interneurons, the AIS diameter was larger in glutamatergic pyramidal cells than in inhibitory neurons (Figs. 8G and 8H).

To complement these results, we also examined the expression of MAP2 and AnkG proteins in neuronal cultures and cerebral cortex from WT and *Slack-R455H* mutant mice by western blotting analysis. We found that levels of AnkG protein were increased, while MAP2 levels were not altered in protein lysates from *Slack-R455H* mutant cultures (Fig. 8J) or cerebral cortex (Fig. 8K) when compared to WT cultures or animals, consistent with the findings in the immunostaining experiments. Taken together, these results suggest that the increased expression of sodium channels in the *Slack-R455H* mutation is accommodated by increases in the length of the AIS in both glutamatergic and GABAergic neurons.

### Proximity of the AIS to the soma is different in glutamatergic neurons and GABAergic neurons

The different subcellular localizations of Slack channels and Na_V_ isoforms raise the possibility that the proximity of potassium K_Na_ channels to sodium Na_V_ channels could be different in excitatory and inhibitory neurons. To test this possibility, we analyzed the relative distance between the soma and the AIS in wild type and mutant mice (“gap length”). In DIV 14 cultures, we observed that the distance between the soma and the AIS was significantly longer in GABAergic neurons than in glutamatergic neurons, but that this was not altered significantly by the *Slack-R455H* mutation in either cell type (Fig. 8F). Similar findings were also observed in the cerebral cortices of 2-month-old mice, in which the distance of the AIS from the soma (defined as the distance between the soma and beginning of immunostaining against Na_V_1.6, Fig. 8I) was longer in inhibitory neurons than in glutamatergic pyramidal cells but this was not affected by *Slack-R455H* mutation.

To test for a direct interaction of Slack channel with Na_V_ subunits, we carried out experiments on mouse cerebral cortex to immunoprecipitate Slack channels, and test for co-immunoprecipitation of Na_V_1.1, Na_V_1.2, and Na_V_1.6 antibodies by western blotting. Our previous studies showed that Slack channel interacts with a protein network including FMRP (Fragile-X Mental Retardation Protein), CYFIP1 (Cytoplasmic FMRP-Interacting Protein-1) and Phactr-1 (Phosphatase and Actin Regulator 1) in its C-terminal domain. Using FMRP and PP1 (protein phosphatase 1, binding protein of Phactr-1) as positive controls, we found that FMRP and PP1 could be readily precipitated by Slack in mouse cortex. Nevertheless, Na_V_1.1, Na_V_1.2, and Na_V_1.6 could not be detected in the Slack-immunoprecipitate (Fig. 8L), suggesting that Slack channels do not directly interact with Na_V_1.1, Na_V_1.2, or Na_V_1.6 subunits in mouse cortex.

## DISCUSSION

We have found that both excitatory and inhibitory neurons of the cerebral cortex of animals expressing a disease-causing mutation in the Slack potassium channel have increased K_Na_ currents and voltage-dependent Na^+^current. Although the gain-of-function Slack mutation did not alter levels of Slack protein either in cultures of cortical neurons or in cerebral cortices of mutant animals, it substantially increased levels of the sodium channel subunit Na_V_1.6, and, in the neuronal cultures, of Na_V_1.1, Na_V_1.2. The mutation-induced increase in sodium channels was associated with longer axonal initial segments and with enhanced expression of ankyrin G, a key scaffold protein of the AIS. A similar increase in Na_V_1.6 staining at the AIS, coupled to increased excitability of pyramidal neurons, had been found in a rat model of temporal lobe epilepsy ^29^.

Previous studies have demonstrated that disease-causing mutations in Slack channel, particularly the MMPSI mutation *Slack-R455H*, markedly increases K^+^current in heterologous cells, with no change in levels of protein or RNA stability of the channel ^3,4^. Heterologous systems, however, provide very limited information on the impact of these variants on neuronal physiology and network excitability and cannot address the cellular background, variety, and subcellular specializations of neurons ^30^. The present study has provided direct evidence that cortical neurons expressing the *Slack-R455H* mutation have increased peak K_Na_ current amplitudes, by as much as 2-4-fold in heterozygous neurons and 4-6-fold in homozygous neurons over that in wild type neurons. It has also shown, however, that, in neurons, mutation of the channel produces cellular effects beyond a simple increase in potassium current.

Work with heterologous expression systems has revealed some of the diverse factors that contribute to the increased K^+^current in different Slack GOF mutations. These include enhanced Na^+^sensitivity, increased channel open probability and/or inter-channel cooperativity ^31,32^. In neurons, however, Slack channels can be activated by Na^+^entry through Na_V_ channels ^33^, non-selective cation channels ^34^or ionotropic receptors such as AMPA receptors ^35,36^, and changes in any of these pathways will also influence K_Na_ currents. Thus, in addition to the increased intrinsic single channel activity of *Slack-R455H* mutant channels ^31^, the increase in I_Na_ produced by the mutation in neurons provides another mechanism for increased K_Na_ current.

The mechanism by which a gain-of-function in Slack channels increase Na_V_ channel expression has not yet been established. One hypothesis, however, is that Slack channel activity directly stimulates the translation of mRNAs encoding Na_V_ subunits. The cytoplasmic C-terminus of Slack channels binds FMRP and CYFIP1 ^37-39^, two established regulators of translation, and the channel co-immunoprecipitates mRNA targets of FMRP ^37^, one of which is mRNA for Na_V_1.6 ^40^. Slack channels are required for changes in intrinsic excitability that require new protein synthesis ^38^and ongoing experiments have found that activation of WT Slack channels stimulates translation of β-actin, another FMRP target ^41^. Future studies will clearly be needed to test the hypothesis that Slack activation directly stimulates the synthesis of Na_V_ subunits.

Paradoxically, we found that the excitability of excitatory neurons was enhanced by the *Slack-R455H* mutation but that of inhibitory interneurons was suppressed. The voltage-dependence of the increased K_Na_ current was different in GABAergic and glutamatergic neurons. In GABAergic neurons, significant increases in K_Na_ current were detected at more negative membrane potentials than in glutamatergic neurons, suggesting that the *Slack-R455H* mutation modifies the intrinsic excitability of GABAergic neurons near the resting potential, hyperpolarizing the neurons and reducing excitability ^8,9,12,13^. Conversely in glutamatergic neurons of *Slack-R455H*, the increased K_Na_ current activates selectively at higher membrane potentials such as those reached only during action potentials, shortening the action potentials and limiting Na^+^channel inactivation. This effect of increased K_Na_ currents to increase excitability has also been found in human IPS cell-derived neurons expressing another Slack channel mutation ^5^.

There are two potential explanations for the opposite effects of increased K_Na_ currents on the excitability of excitatory and inhibitory neurons. The first is that excitatory and inhibitory cortical neurons may express different splice isoforms of KCNT1 channel. Two splice isoforms, Slack-A and Slack-B, have been characterized, and these differ only at their N-terminal cytoplasmic domain ^42^. Slack-A isoform activates and deactivates very rapidly with voltage. Such rapid activation during an action potential is known to increase the rate of repolarization and subsequent rapid afterhyperpolarization. In contrast, Slack-B isoform activates more slowly and at more negative potentials ^42^. An increase in such a slowly activating K^+^current at the resting membrane potential would increase the amount of current needed to evoke action potentials and reduce excitability. It is possible that excitatory neurons primarily express the rapidly-activating Slack-A isoform while inhibitory neurons express the slowly-activating B-splice isoform channels.

Independently of which splice isoform is expressed, a second factor that could determine whether increased K_Na_ currents promote or suppress hyperexcitability is the proximity of the K_Na_ channels to the source of Na^+^influx. Patch clamp recordings of clusters of K_Na_ channels at the nodes of myelinated axons suggest that the rise in Na^+^concentration during a single action potential is sufficient to activate these channels ^43^. Rapid, transient and very local activation of K_Na_ channels by a large Na^+^transient during an action potential would be expected primarily to produce rapid and transient repolarization, enhancing excitability with little effect on resting membrane properties, as we observed for excitatory neurons (Fig.3). In contrast, slower K_Na_ activation by the accumulation of Na^+^from an influx source remote from the K_Na_ channel would be expected to influence resting membrane properties by increasing input resistance and suppressing repetitive firing, as we observed in inhibitory neurons (Fig. 4).

While the light level immunocytochemistry we carried out on neurons from wild type and mutant animals does not have the resolution to differentiate different degrees of colocalization or clustering of Slack channels with Na_V_ channels at the molecular level, it revealed clear difference in the organization of the AIS in excitatory neurons and inhibitory neurons. The AIS plays a key role in regulating neuronal excitability ^44,45^, and previous work has indicated that the organization of the AIS is heterogenous in cortical neurons ^46^. Our studies on frontal cortex found that the distance between the soma and the AIS was significantly longer in GABAergic interneurons than in pyramidal neurons. Because Slack staining is abundant at the soma, this provides another potential mechanism for differences in activation of K_Na_ channels in the two types of cells. Although we were not able to detect coimmunoprecipitation of Slack with any of the Na_V_ subunits, which would support tight colocalization at the molecular level, our work is consistent with previous studies showing that Na_V_1.1, Na_V_1.2 and Na_V_1.6 have distinct cell type specificities and subcellular distributions. Specifically, Na_V_1.2 and Na_V_1.6 are highly expressed at the AIS and nodes of Ranvier of excitatory pyramidal neurons, while Na_V_1.1 is more associated with inhibitory interneuron ^25-28^. We also found that Na_V_1.2 and Na_V_1.6 were highly expressed at the AIS of cultured cortical neurons, and that Na_V_1.1 was expressed at the AIS as well as at the soma and dendrites. Further studies of the chemical and spatial interactions between Slack channels and channels that provide Na^+^ influx will be required to resolve some of these issues and provide new therapeutic directions for the treatment of Slack-related epilepsies and intellectual disability.

## ACKNOWLEDGMENTS

This research was supported by NIH Grants NS102239 to LKK.

## AUTHOR CONTRIBUTIONS

Conceptualization, J.W. and L.K.K.; Methodology, J.W., I.R.Q., Y.L.Z., and L.K.K.; Formal Analysis, J.W., I.R.Q., Y.L.Z., M.B., and L.K.K.; Investigation, J.W., I.R.Q., Y.L.Z., AND M.B.; Writing – Original Draft, J.W. and L.K.K.; Writing – Review & Editing, J.W. and L.K.K.; Supervision, L.K.K.; Funding Acquisition, L.K.K.

## Supplementary Information

**Figure S1. TTX-sensitive K**^**+**^**currents are increased in cortical neurons expressing the *Slack-R455H* mutation. A**. Representative current traces from whole cell voltage clamp recordings in physiological extracellular medium without TTX (*left*) and with 0.5 μM TTX (*right*) in response to voltage steps (−90 to +50 mV) in WT (*black*), *Slack*^*+/R455H*^ (*blue*), and *Slack*^*R455H/R455H*^ (*red*) neurons. **B**. Summary data show the TTX-sensitive K^+^current, which was calculated by subtracting the trace obtained with TTX from the trace without TTX, for each voltage step in primary cortical neuron. Data are shown as mean ± SEM (n = 8-10, two-way ANOVA). **C**. Reduction of K^+^current at +50 mV for each genotype and neuron type. Data are shown as mean ± SEM (n = 8-10, one-way ANOVA). \**p <* 0.05, \*\**p <* 0.01, \*\*\**p <* 0.001 and \*\*\*\**p <* 0.0001.

**Figure S2. Electrophysiological parameters of current clamp recording from primary cortical neurons. A-D**. RMP (**A**), AP threshold (**B**), depolarization rate (**C**), and repolarization (**D**) of recorded glutamatergic neurons from WT (*black*), *Slack*^*+/R455H*^ (*blue*), and *Slack*^*R455H/R455H*^ (*red*) littermates. **E-H**. RMP (**E**), AP threshold (**F**), depolarization rate (**G**), and repolarization (**H**) of recorded FS-GABAergic neurons from each genotype. **I-L**. RMP (**I**), AP threshold (**J**), depolarization rate (**K**), and repolarization (**L**) of recorded NFS-GABAergic neurons from each genotype. Data are shown as mean ± SEM (n = 12-20, one-way ANOVA). \**p <* 0.05 and \*\**p <* 0.01.

**Figure S3. Subcellular localizations of Na**_**V**_**1.1 and Na**_**V**_**1.2 channels in primary cortical neuron. A**. Immunostaining of Na_V_1.1 with dendrite (MAP2) and AIS (AnkG) markers in cultured cortical neurons on DIV 14. Glutamatergic and GABAergic neurons were differentiated according to their morphological features. Scale bar, 50 μm. **B. M**Immunostaining of Na_V_1.2 with dendrite (MAP2) and AIS (AnkG) markers in cultured cortical neurons on DIV 14. Glutamatergic and GABAergic neurons were differentiated according to their morphological features. Scale bar, 50 μm.

**Figure S4. Subcellular localizations of Na**_**V**_**1.1 and Na**_**V**_**1.2 channels in cerebral cortex**.

**A**. Immunostaining of Na_V_1.1 with AnkG (top, Scale bar, 50 μm) and GAD67 (bottom, Scale bar, 25 μm) in cerebral cortex of 2-month-old mice. **B**. Immunostaining of Na_V_1.2 with AnkG (top, Scale bar, 50 μm) and vGLUT1 GAD67 (bottom, Scale bar, 25 μm) in cerebral cortex of 2-month-old mice. Glutamatergic and GABAergic neurons were differentiated by their co-immunostaining with vGLUT1 and GAD67. **C**. Representative images of a single neuron showing the possible colocalization areas of Slack and Na_V_1.6. Scale bar, 10 μm.

### METHODS

**KEY RESOURCES TABLE**

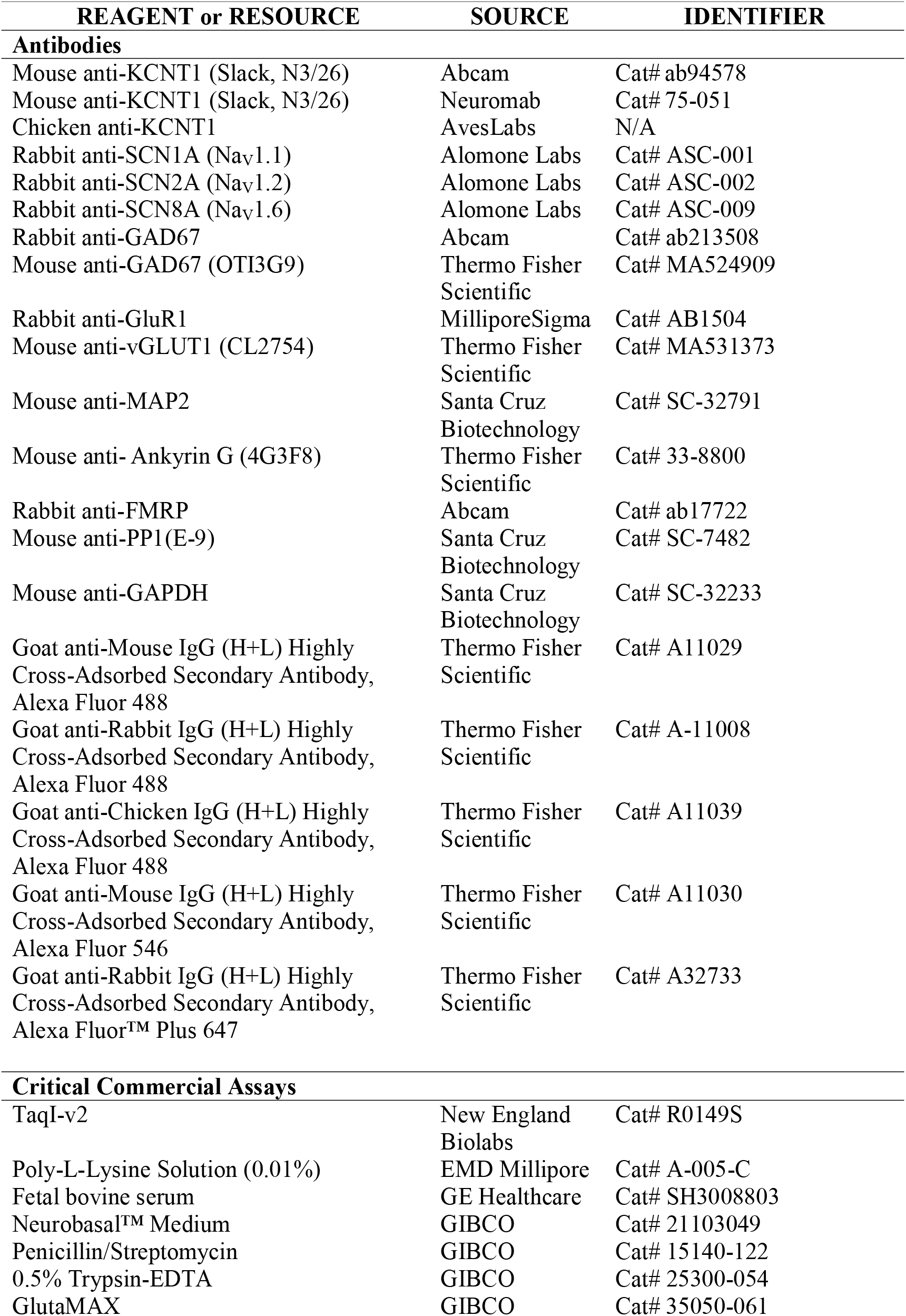
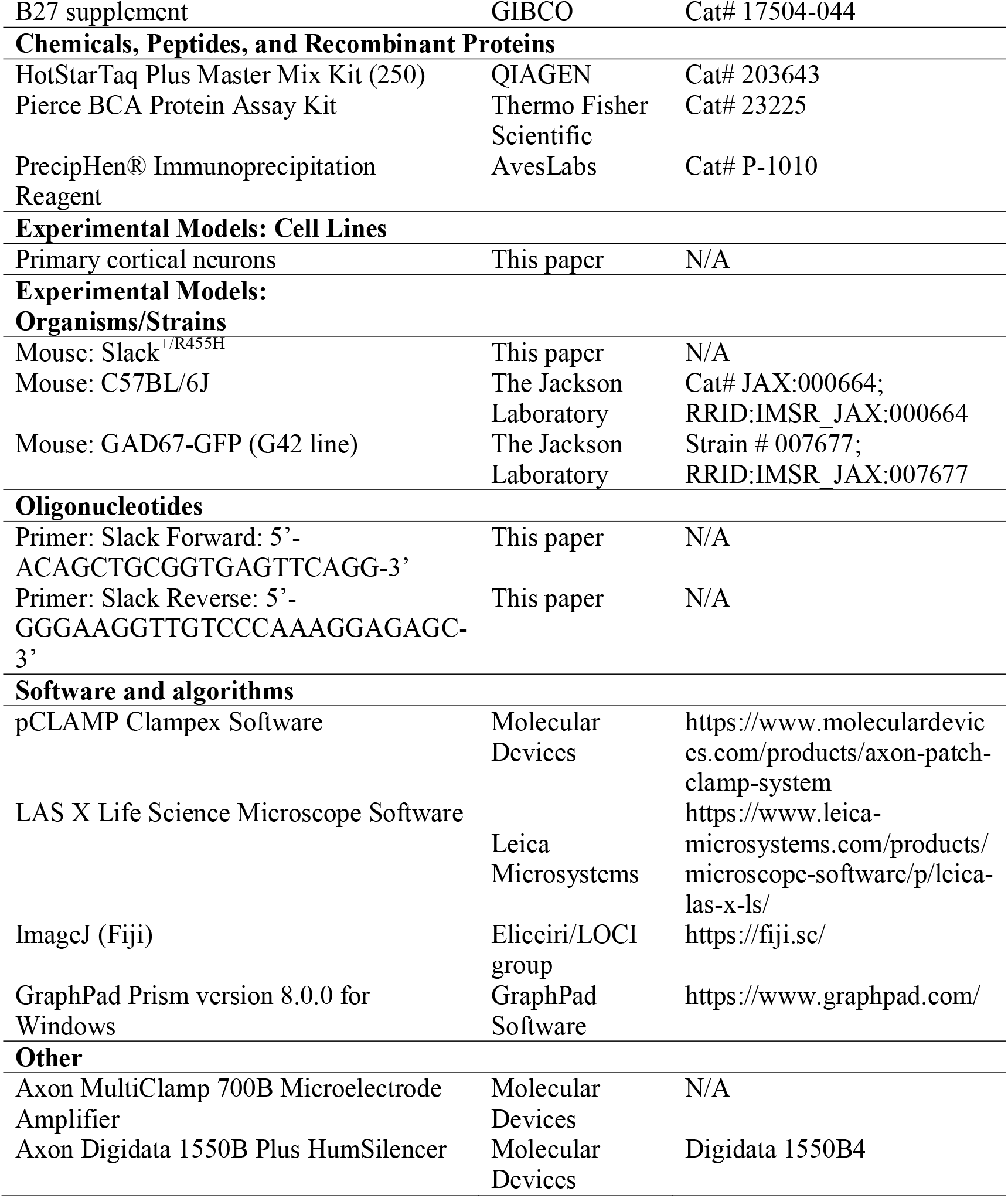

#### RESOURCE AVAILABILITY

##### Lead Contact

Further information and requests for resources and reagents should be directed to and will be fulfilled by the Lead Contact, Leonard K. Kaczmarek.

##### Materials Availability

The *Slack*^*+/R455H*^ mouse line is available from the Lead Contact, Leonard K. Kaczmarek, with a completed Materials Transfer Agreement.

##### Data and Code Availability

The published article includes all datasets generated or analyzed during this study. Any additional information required to reanalyze the data reported in this paper is available from the Lead Contact upon request.

#### EXPERIMENTAL MODEL AND SUBJECT DETAILS

##### Mouse Strains

*Slack*^*+/R455H*^ mice were generated in the C57BL/6J mouse strain using CRISPR/Cas9 via the Yale Genome Editing Center. The generation and basic characterization of this mouse model are available in our previous paper ^18^. Heterozygous (WT/R455H, *Slack*^*+/R455H*^) mice were used as breeding pairs to obtain WT, heterozygous and homozygous (R455H/R455H, *Slack*^*R455H/R455H*^) embryos for study. Littermates were labeled and genotyped using gene-specific polymerase chain reaction (PCR) on DNA extracted from tail tissues with primers (*Slack* forward primer: 5 ‘-ACAGCTGCGGTGAGTTCAGG-3 ‘; *Slack* reverse primer: 5 ‘-GGGAAGGTTGTCCCAAAGGAGAGC-3 ‘) with standard thermocycler amplification conditions. Following amplification, a restriction cut was performed with the enzyme TaqI-v2 (NEB, R0149S) to distinguish WT (196 and 112 bp products after cut), R455H heterozygous (112, 196, and 308 bp products) and homozygous (308 bp product) alleles. Whenever possible, investigators were blind to the genotype of the mice. GAD67-GFP mice were purchased from Jackson Laboratories (Bar Harbor, MA). For all experiments, male and female littermates were used for each genotype. All experiments were performed in accord with the NIH Guidelines for the Care and Use of Laboratory Animals and approved by Yale University ‘s Institutional Animal Care and Use Committee (IACUC).

##### Primary Cortical Neuron Culture

Primary cortical neurons were prepared from E16-17 mouse embryos as described previously with modifications specific for this study ^47^. After isolation of frontal cortex from embryonic brains, neurons were dissociated and seeded (on coverslips inside a 6 well plate: 0.2E6 cells/well) onto plates containing NB plus [Neurobasal medium supplemented with B27 (Invitrogen GIBCO Life Technologies), GlutaMAX (GIBCO), and penicillin/streptomycin (GIBCO)] and 5% FBS (GIBCO). After 2 hours incubation, primary cultures were maintained in NB plus without FBS in a 5% CO_2_ and 20% O_2_ incubator at 37 °C. Subsequently, half the medium was replaced every 2 days.

### METHOD DETAILS

#### Immunofluorescence

For immunocytochemistry of primary cortical neurons on DIV 14, coverslips were fixed in 4% paraformaldehyde for 10□min and permeabilized in blocking buffer (0.2% Triton X-100 and 3% normal goat serum in PBS) for 5□min at room temperature. For immunohistochemistry of cerebral cortex from two-month-old mice, brains were sectioned into 100-mm coronal slices on a vibrating microtome (Leica) and post-fixed in 4% PFA overnight at 4 °C. Brain slices were then permeabilized and incubated in blocking buffer for one hour at room temperature. After blocking, sections from DIV 14 cortical neuronal cultures or two-month-old mice were overlaid with primary antibodies overnight at 4 °C. Then, the corresponding Alexa Fluor 488-, 546- or 647-conjugated secondary antibodies were applied. Stained sections were mounted with DAPI-containing mounting solution and sealed with glass coverslips.

Confocal imaging of primary cortical neurons and brain slices was carried out on a Leica SP5 MP. For AIS analysis, three-dimensional z-stack images were taken and stitched into tiles to cover the entirety of the neuron of interest. Z-stacks were then loaded in Leica LAS-X. AIS were defined as AnkG/Na_V_1.6-labeled segments with clearly identifiable start and end points. AIS start point was defined as sharply increased AnkG/Na_V_1.6 signal closest to the soma. AIS end point was defined as reduced AnkG/Na_V_1.6 signal to the point where it could no longer be discerned from the background. AIS that had blunt ends were excluded, as this is likely an artifact of cutting through the AIS during sectioning. The AIS length measurement was performed by carefully tracing the shape of the AIS using segmented line tool in Leica LAS-X. The length was measured in around 3 AIS per image, and three images were analyzed per mouse or culture. Measurement of distance from soma to AIS start point was performed as described previously ^25, 48, 49^.

#### Western blot and Co-immunoprecipitation (Co-IP)

Protein lysates from DIV 14 cortical neurons and two-month-old mice frontal cortex were prepared using Pierce IP Lysis Buffer (Thermo Scientific) supplemented with cOmplete EDTA-free Protease Inhibitor Cocktail (Millipore Sigma) according to manufacturer ‘s protocol. Protein quantification was performed using Pierce BCA Protein Assay Kit (Thermo Scientific).

For Co-IP experiments, cerebral cortex lysates from WT mice were incubated with 5μg anti-Slack IgY antibody or IgY control (AvesLabs) overnight at 4°C. 100 μL Anti-IgY PrecipHen beads (AvesLabs) was added to sample and allowed to incubate for 2 hours, followed by wash and collection of beads. Beads were transferred to 2x Laemelli-Buffer with 5% beta-mercaptoethanol and incubated at room temperature for 30 minutes prior to Western blotting ^39^. For all immunoblotting experiments, protein samples were electrophoretically separated on an SDS-PAGE gel (4%–15% gradient gel, Bio-Rad) and transferred onto PVDF membranes (0.2 μm pores, Bio-Rad, USA). Blots were blocked in 5% nonfat milk in Tris-buffered saline and Tween 20 (TBST) for 1 h at room temperature and probed with the primary antibody in 5% milk-TSBT overnight at 4°C. After overnight incubation, the blots were washed three times in TBST for 30min, followed by incubation with corresponding horseradish peroxidase-conjugated secondary antibodies (1:1000; Abcam, UK) at room temperature for 1 h. Protein bands were visualized via enhanced chemiluminescence and quantified with analyzed with ImageJ (NIH) software. GAPDH was used as a loading control levels.

#### Patch-clamp Recordings

Whole-cell patch-clamp recordings were performed with patch-clamp amplifiers (MultiClamp 700B; Molecular Devices) under the control of pClamp 11 software (Molecular Devices). Data were recorded with a sampling rate at 20 kHz and filtered at 6 kHz. Rs compensation of 70% was used. Primary cortical neurons at DIV 13-14 were recorded at physiological temperature (37°C). Recording electrodes were pulled from filamented borosilicate glass pipettes (Sutter Instrument, CA), and had tip resistances between 4 and 6 M? when filled with the following internal solution (in mM): 124 K-gluconate, 2 MgCl_2_, 13.2 NaCl, 1 EGTA, 10 HEPES, 4 Mg-ATP, and 0.3 Na-GTP (pH 7.3, 290-300 mOsm). The extracellular medium contained the following (in mM): either 140 NaCl or 140 N-methyl-D-glucamine (NMDG), 5.4 KCl, 10 HEPES, 10 glucose, 1 MgCl_2_, and 1 CaCl_2_ (pH 7.4, 310 mOsm). K_Na_ currents and TTX-sensitive K^+^ currents were recorded according to the protocols reported in previous studies ^5,13^. Neurons were held at -80 mV and given 60 ms voltage pulses in 10 mV steps over a range of -90 to +50 mV. The difference current over the 20 ms at the end of the voltage pulse was considered the steady state K_Na_ current. To isolate the K_Na_ currents, whole-cell voltage recordings were performed using physiological extracellular medium with 140 mM Na^+^ ions and without Na^+^ (140 mM NMDG). The traces obtained without Na^+^ were then subtracted from the traces obtained in the external Na^+^ medium ^5^. To isolate the TTX-sensitive K^+^ currents, current traces from the TTX (0.5 μM) solution were subtracted from the current traces obtained from the standard solution to obtain the difference current ^12^.

For current-clamp recordings, the firing properties of neurons were tested by injecting 200 ms square current pulses incrementing in 20 pA steps, starting at -20 pA. To compare the maximum number of action potentials (APs), 1.5 s square current pulses in 20 pA steps were injected until the number of APs per stimulus reached a plateau phase. The AP, amplitude, half width, AHP, RMP, threshold, depolarization rate and repolarization rate values, as well as input resistance and rheobase were analyzed according to the protocols reported in previous studies ^5,13^ and using the Clampfit 11 inbuilt statistics measurements program.

#### QUANTIFICATION AND STATISTICAL ANALYSIS

Normality and variance similarity were measured by GraphPad Prism before we applied any parametric tests. Two-tailed Student ‘s t test (parametric) or unpaired two-tailed Mann-Whitney U-test (non-parametric) was used for single comparisons between two groups. Other data were analyzed using one-way or two-way ANOVA with Bonferroni correction (parametric) or Kruskal-Wallis with Dunn ‘s multi comparison correction (non-parametric) depending on the appropriate design. Post hoc comparisons were carried out only when the primary measure showed statistical significance. All data were expressed as mean ± SEM, with statistical significance determined at p values < 0.05. In details, *Indicates *p <* 0.05; \*\**p <* 0.01; \*\*\**p <* 0.001; \*\*\*\**p <* 0.0001 in all figures.

