## Supplementary figures and images for "Disease-causing Slack potassium channel mutations produce opposite effects on excitability of excitatory and inhibitory neurons"

### Supplemental Figure 1

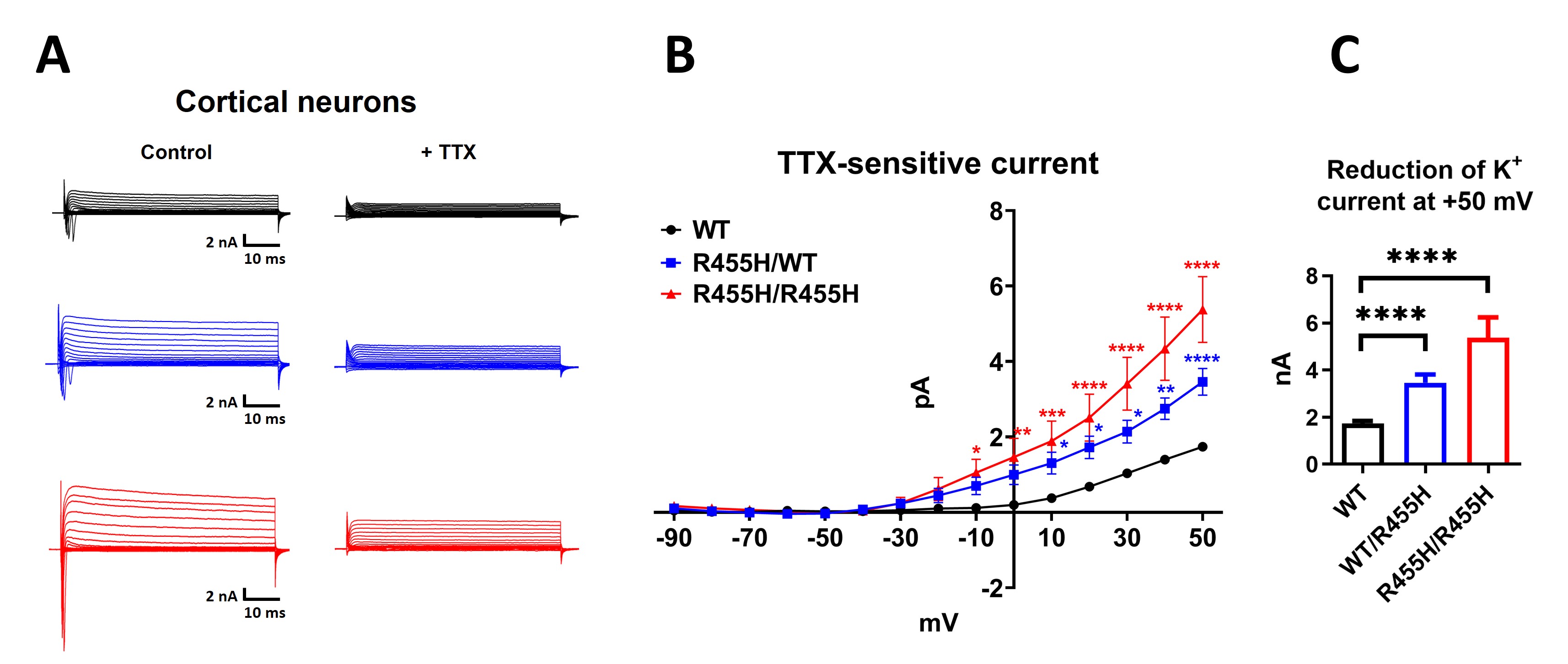

### Supplemental Figure 2

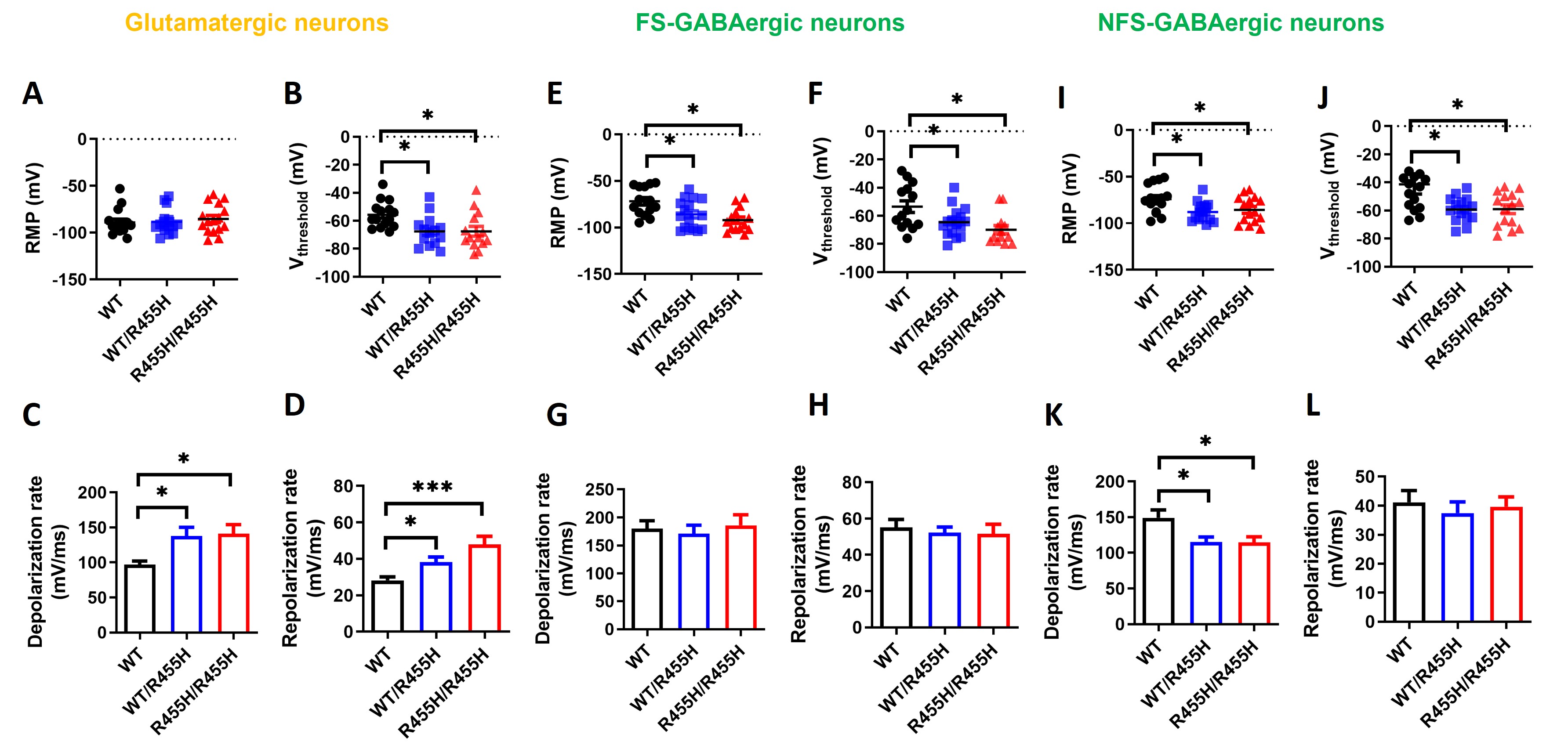

### Supplemental Figure 3

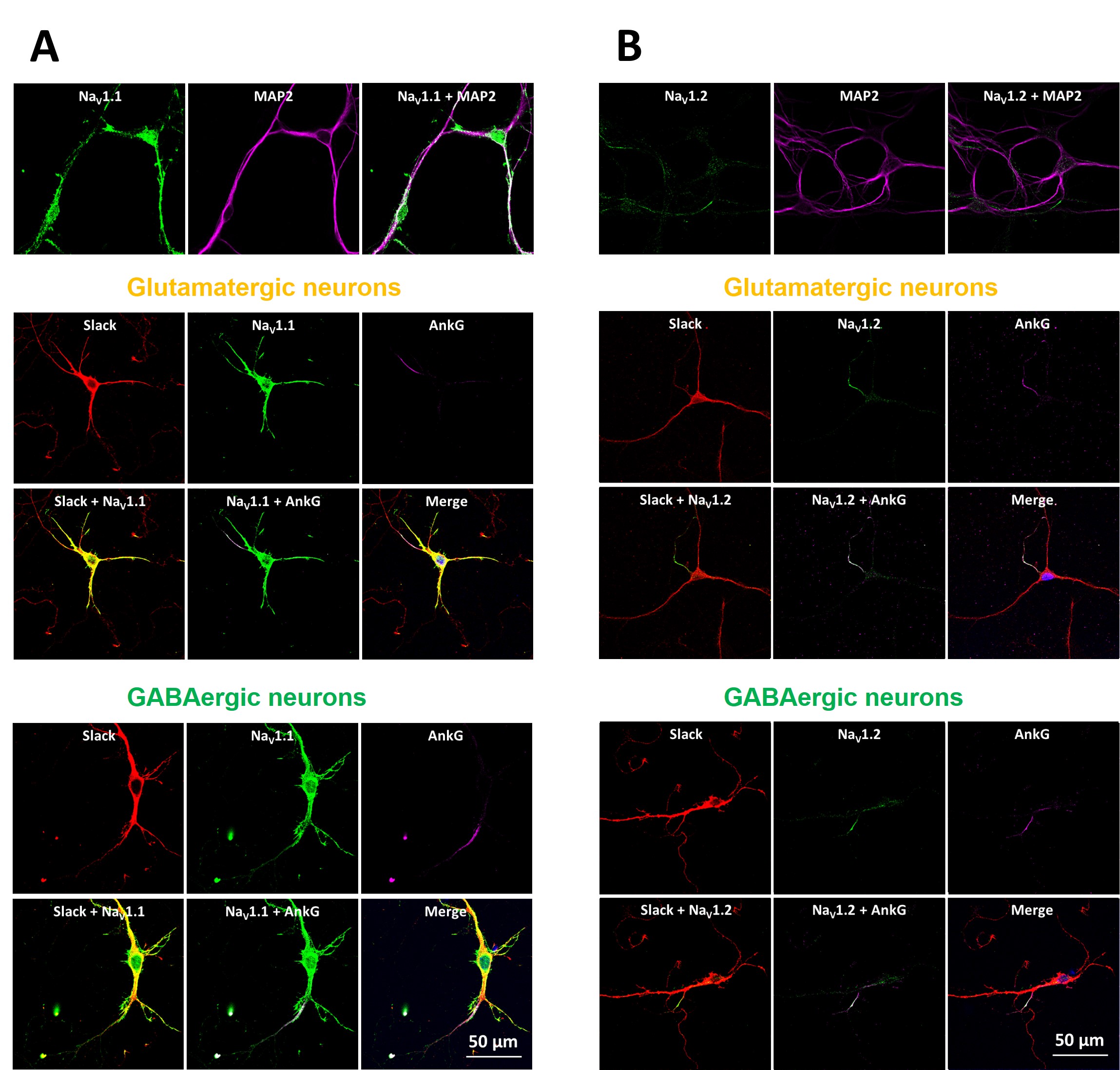

### Supplemental Figure 4

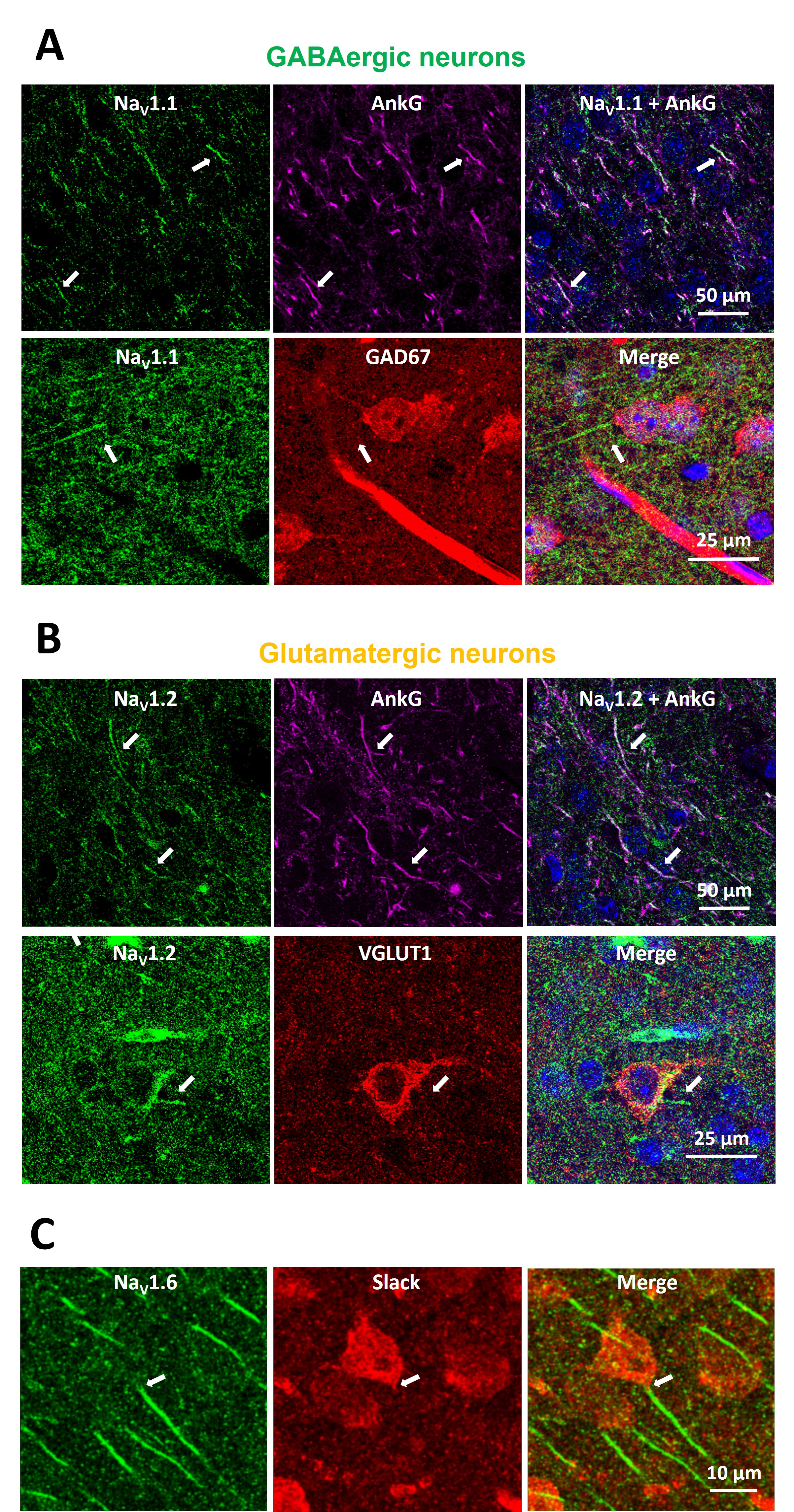
